# Whole-genome sequencing of Bacillus thuringiensis strain SY49.1 reveals the detection of novel candidate pesticidal and bioactive compounds isolated from Turkey

**DOI:** 10.1101/2022.03.07.482483

**Authors:** Semih Yılmaz, Abeer Babiker Idris, Abdurrahman Ayvaz, Rıdvan Temizgül, Mohammed A. Hassan

**Author notes:** Authors’ E-mails: SY; ABI; AA; MAH.

## Abstract

**Background:** *Bacillus thuringiensis* SY49.1 *(Bt* SY49.1) strain is a new strain isolated from a soil sample in Adana city which is nestled in the most fertile agricultural area in Turkey. This strain has insecticidal activity against insect pests from different orders. Also, it is characterized by its significant activity against plant fungal infections and as plant growth promotion.

**Aims:** To report the whole genome sequencing (WGS) and annotation of the *Bt* SY49.1 strain.

**Materials and Methods:** The *Bt* SY49.1 strain was isolated from the soil sample in Adana city by using a sodium acetate enriched medium. Bacterial DNA was extracted then sequenced using Illumina HiSeq technology. After data filtering and quality control, de novo assembly and genomic annotation were applied. Evolutionary and comparative genomic analysis and variant calling were performed using various in silico tools.

**Results:** The WGS of strain *Bt* SY49.1 is 6. 32 Mbp long with a GC content of 34.68%. It reveals large numbers of single nucleotide polymorphisms (SNPs) and InDels. The assembled genome contains 6,562 protein-encoding genes. In addition, it encodes various gene inventories for the biosynthesis of bioactive compounds such as insecticidal delta endotoxins, growth stimulatory deaminase and phosphatases, fungicidal thuricin, bacillibactin, petrobactin, fengycin / plipastin, and zwittermycin A.

**Conclusion:** The strain *Bt* SY49.1 could have several potential utilities as a source of antibiotics compounds, plant growth promoting metabolites, and biocontrol agents for fungal phytopathogens, and insects. We expect that the draft genome of the *Bt* SY49.1 strain may provide a model for proper understanding and studying of antimicrobial compound mining, genetic diversity among the *B. cereus* group, and pathogenicity against insect pests and plant diseases, and growth-promoting activity.

## 1. Introduction

*Bacillus thuringiensis* (*Bt*) is a ubiquitous rod-shaped, Gram-positive, spore-forming and aerobic or facultative anaerobic bacterium found usually in a variety of ecological niches including soil, grain dust, dead insect, and aquatic environments, among many others (1, 2). It is characterized by producing parasporal crystals, during its sporulation or stationary growth phase (3). These crystals contain protein toxins (e.g. δ-endotoxins or Cry proteins) which act as powerful intestinal toxins against various insect hosts (4, 5). The toxins are active against larvae of lepidopteran, dipteran, and /or coleopteran species and have been successfully introduced to genetically modified crops. Their lack of toxicity against mammals, high specificity, and environmental safety led to advancing in search for new strains with different spectra of pesticide activities (4, 6). In addition, *Bt* strains have been found to exhibit antibacterial, antibiofilm, antifungal, and emulsifying activities (7-9). They are known to be significant sources for antimicrobial compounds, such as bacteriocins and lipopeptides, and antifungal compounds, such as zwittermycin, lipopeptides, and chitinase (8, 10, 11).

Several strains of *B. thuringiensis* have raised a global interest for various bio-pest applications because of their specific pesticidal activities (12-16). However, the development of insect resistance to *B. thuringiensis* toxin proteins has been widely reported (17, 18). This leads to urgent searching for novel *B. thuringiensis* with novel insecticide toxins as an effective management strategy (19). In this study, we report the complete genome sequence of a novel *B. thuringiensis* SY49.1 strain which was isolated from a soil sample collected from Adana city in Turkey. This strain has insecticidal activity against insect pests from different orders; and was found to carry various types of insecticidal *cry* genes which include *cry1Aa/Ad, cry1B, cry1C, cry5, cry9A*, and *cry9C* (20-23). Also, it is characterized by its significant activity against plant fungal infections (24).

## 2. Materials and Methods

### 2.1 Bacterial isolation and identification

*Bt* SY49.1 strain was isolated from a soil sample collected from Adana city which is nestled in the most fertile agricultural area in the south of Turkey. The isolation of the *Bt* SY49.1 strain was done by using a sodium acetate enriched medium, as previously described (25). In brief, one gram of soil sample was inoculated in LB broth (pH 6.8±2) which includes 0.25M sodium acetate, then incubated in a shaking incubator at 200 rpm at 30°C for 4 hours. After that, 1.5 ml of the inoculated LB broth has been transferred to a sterile Eppendorf tube and exposed to high temperature (80°C) for 10 minutes to kill the vegetative bacterial form. Then, 20-50 ul from the Eppendorf tube were spread on LB agar plates and incubated overnight at 30°C. Finally, colonies with a morphological appearance that resembles *Bacillus* have been spread on agar plates to obtain pure colonies which were re-incubated in 5 ml LB broth (pH 6.8±2) in 50 ml tubes at 200 rpm at 30°C overnight. Further bacterial identifications were performed by colonial morphology on Potato Dextrose Agar (PDA), gram stain, and malachite green stain as previously described in (26), (27), and (28), respectively.

For para-sporal inclusion bodies characterization, the bacteria were inoculated in 150 ml of 3T medium and incubated at 30 °C for 7 days to induce sporulation (20). After that, the mixtures were centrifuged at 15.000 rpm and 4 °C for 10 min and suspended in dH_2_O on microscope slides and fixed. Then, the slides were sputter-coated with 10 150 nm Au / Pd using an SC7620 Mini-sputter coater. Finally, they were viewed using a LEO440 scanning electron microscope at 20kV beam current (29).

### 2.2 Bacterial DNA extraction and sequencing

The bacterial DNA was extracted according to a previously described method of Jensen *et al*. and Porcar *et al*. with some modifications (30, 31). In brief, a loopful of cells from a single colony, that has been cultured in LB medium overnight, was placed into 400 μl sterile dH_2_O. Then, to lyse the cells, the mixture was boiled for 10 min. After that, the resulting cell lysate was centrifuged for 10 sec at 10.000 rpm. Finally, the supernatant was used as DNA templates for PCR reactions.

The whole-genome sequencing (WGS) was commercially conducted using Illumina’s HiSeq technology at BGI-HongKong Co., Ltd. company.

### 2.3 Genome assembly and genome annotation

A 760.2 Mbp paired-end reads with a 500-bp insert size library were obtained from the DNA sample. However, to ensure accuracy of the follow-up analysis 10-20% of the data was eliminated after filtering the raw data and removing reads with low-quality, reads with a certain proportion of Ns’ bases, adapter contamination, and duplication reads. For K-mer analysis (32), regions in the reads with high-quality scores were intercepted and, then, cut into 15-bp fragments (15-mer). The depth of 15-mer was 16x and the genome coverage depth was 18.96x. In addition, by using SOAPaligner (version 2.21) software (33), all read sequences obtained by sequencing were initially aligned with the reference sequence (NC_017208) to calculate the average depth and coverage ratio in order to determine the differences between sequencing species and the reference sequence. The Coverage depth and coverage ratio of *Bt* SY49.1 strain to reference sequence (*Bt* chinensis-CT-43 serovar) were 48.79x and 80.29%, respectively.

De novo assembly to the short filtered reads was performed using SOAPdenovo (version 1.05) (34), which is an in-house assembler. De Bruijn graph was constructed, firstly, and erroneous connections were removed to build the contig and then scaffold. For filling gaps, reads are mapped to scaffold and a paired-end reads relationship was used. In addition, the assembly error was corrected by aligning the reads to the assembly result using SOAPaligner (version 2.21) tool and counting the mapping information of reads. Then the single base error of the assembly result was corrected based on mapping information. Also, GC content and depth correlative analysis, and K-mer analysis were performed to evaluate whether heterogeneous sequences exist in the sample or not.

Genome annotation was done using the NCBI Prokaryotic Genome Annotation Pipeline (PGAP) and Rapid Prokaryotic Genome Annotation (Prokka), and gene functions were predicted using Rapid Annotation using Subsystems Technology (RAST) and the Pathosystems Resource Integration Center (PATRIC) (35-38). A circular map was generated using PATRIC sever (38).

In addition, CRISPR repeats were predicted by using the CRISPRfinder web server (39). AntiSMASH version 4.0 server was used to predict, annotate, and analyze secondary metabolite biosynthesis gene clusters in the bacterial genome (40). PHASTER web server is used for the identification and annotation of prophage sequences within *Bt* SY49.1 genome and plasmids (41). Also, IslandViewer 4 webserver was applied to predict and interactively visualize genomic islands (GI) in *Bt* SY49.1 genome (42).

### 2.4 Evolutionary and comparative genomic analyses of *Bt* SY49.1 strain

The species was established using *16S rRNA* and phylogenic approach. Sequences of *16S rRNA* genes of thirteen *Bacillus* strains and SY49.1 were selected for phylogenetic analysis. The selected species represent the members of the *B. cereus sensu lato* group. The sequence of the strain *B. subtilis* IAM 12118 was selected as an outgroup. The Kimura 2-parameter (K2+G) model from the substitution (ML) model was chosen with 1000 bootstrap replicates to construct a maximum likelihood (ML) phylogenetic tree using MEGA7 software (43). Also, phylogenetic analysis of the strain SY49.1 with the closest reference and representative genomes was conducted using PATRIC sever (38). The closest reference and representative genomes to SY49.1 were determined by Mash /MinHash (44). PATRIC global protein families (PGFams) were selected from these genomes to identify the phylogenetic placement of *Bt* SY49.1 genome (45). Moreover, the strain was established using the Genome-to-Genome Distance Calculator (GGDC) webserver (46).

Functional comparison (sequence-based) of *Bt* SY49.1 genome was performed with closely related *Bacillus* species, *B. thuringiensis* str. 97-27 (NC_005957), *B. thuringiensis* str. Al Hakam (NC_008600), *B. cereus* ATCC 10987 (NC_003909), *B. cereus* ATCC 14579 (NC_004722), *B. cereus* E33L (NC_006274), and *B. anthracis str*. Ames Ancestor (NC_007530). While *B. subtilis* subsp. subtilis str.186 (NC_000964) was used as an outgroup in the map. The comparison was done by using bidirectional and unidirectional best hit (protein Blast) implemented in the RAST server (35). Furthermore, variation analyses for SNP and InDel calling were conducted using MUMmer (version 3.22) package and LASTZ (version 1.01.50) (47, 48).

## 3. Results

### 3.1 Genome properties of strain *Bt* SY49.1

The complete genome sequence of the *Bt* SY49.1 strain is 6,322,230 bp long with a GC content of 34.68%, containing 270 scaffolds with N_50_ of 79,855 bp. The *Bt* SY49.1 genome contains a chromosome and seven circular plasmids bigger than 15kb. The GC contents of the seven plasmids ranged from 31.6% to 36.9%. In total, 6,562 protein-encoding genes (PEGs), 3 *rRNAs*, 10 *tRNAs*, 5 *ncRNAs*, and 429 pseudogenes were annotated by NCBI-PGAP. Two sequences were predicted with CRISPR repeats and one sequence with Cas cluster. The *Bt* SY49.1 whole-genome sequencing project has been deposited at NCBI GenBank under the accession number NZ_JAHKEZ000000000, assembly number GCF_018791885.1 BioProject accession number PRJNA734785, and BioSample accession number SAMN19533886. The 6.32 Mbp draft genome map of *Bt* SY49.1 is presented in Figure 1.

**Figure 1.**
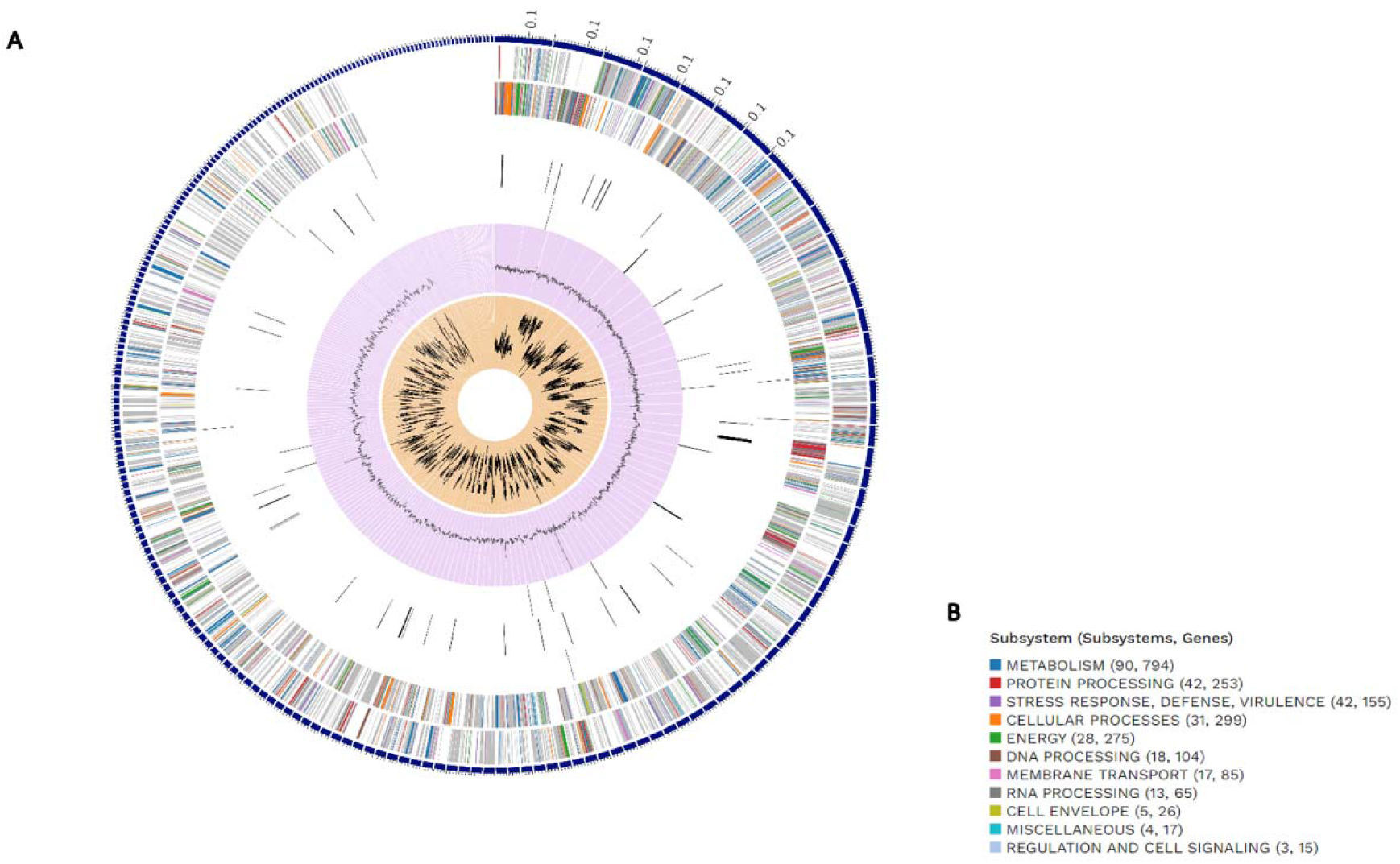
Circular visualization of the draft genome of *Bt* SY49.1 represents relevant genome features. This includes, from outer to inner circles, the contigs, CDS on the forward strand, CDS on the reverse strand, RNA genes, CDS with homology to known antimicrobial resistance genes, CDS with homology to known virulence factors, GC content, and GC skew. The colors of the CDS on the forward and reverse strand indicate the subsystem that these genes belong to, as presented in B). The circular display has been limited to the 230 longest contigs of the 708 contigs in the genome.

### 3.2 Insights from the genome sequence

*Bt* SY49.1 was found to be flagellated, sporulating with a subterminal endospore and producing the insecticidal parasporal inclusions, see Figure 2. These phenotypes are supported by gene inventories observed in the genome of *Bt* SY49.1. The RAST annotation has sorted the 6,562 PEGs into 342 functional subsystem pathways using SEED subsystems. The most abundant of them are genes that are associated with amino acids and derivatives metabolism 392 (18.8%), followed by carbohydrates metabolisms 264 (12.67%), cofactors, pigment, and prosthetic groups 158 (7.59%), protein metabolism 156 (7.5%), and dormancy and sporulation 109 (9. 23%), for more illustration, see Figure 3. In addition, the antiSMASH 4.0 server predicted that the *Bt* SY49.1 genome encodes gene clusters that are responsible for the biosynthesis of secondary metabolites which include lantipeptide, antimicrobial peptides, and siderophores. The *Bt* SY49.1 genome was found to carry gene clusters with high homology to the biosynthetic gene clusters of the thuricin, iron-siderophore (bacillibactin and petrobactin), fengycin / plipastin, and antifungal compound (zwittermycin A) (Figure 4). Moreover, the PHASTER server identified ten prophage regions, of which one region is intact (score > 90), seven regions are incomplete (score < 70), and two regions are questionable (score 70-90), see Figure 5.

**Figure 2.**
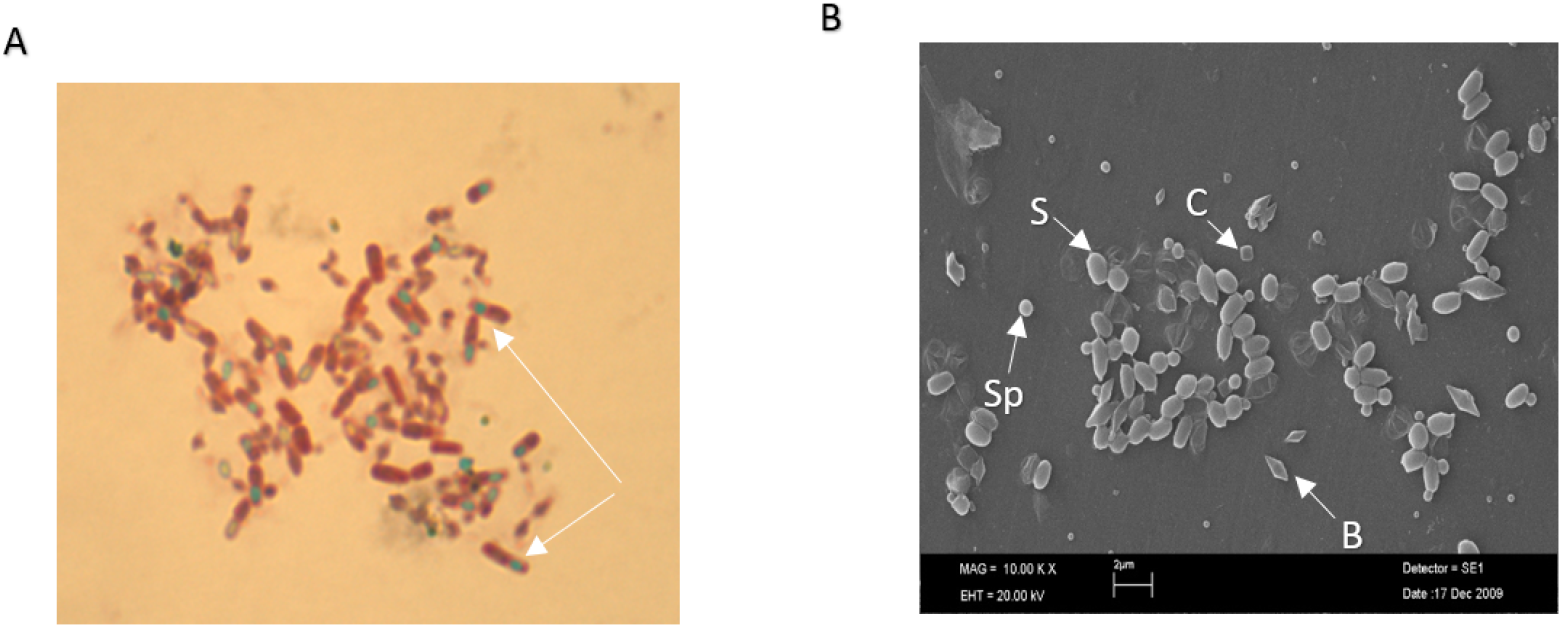
Microscopical characteristic of *Bt* SY49.1. 2A. Microscopic view of *Bt* SY49.1 after spore staining. Arrows indicate the endospores. 2B. Scanning Electron Microscopy (SEM) image of the dfferent types of crystal proteins and spores (S) produced by *Bt* SY49.1. B: bipyramidal, C: cubic, Sp: spherical.

**Figure 3.**
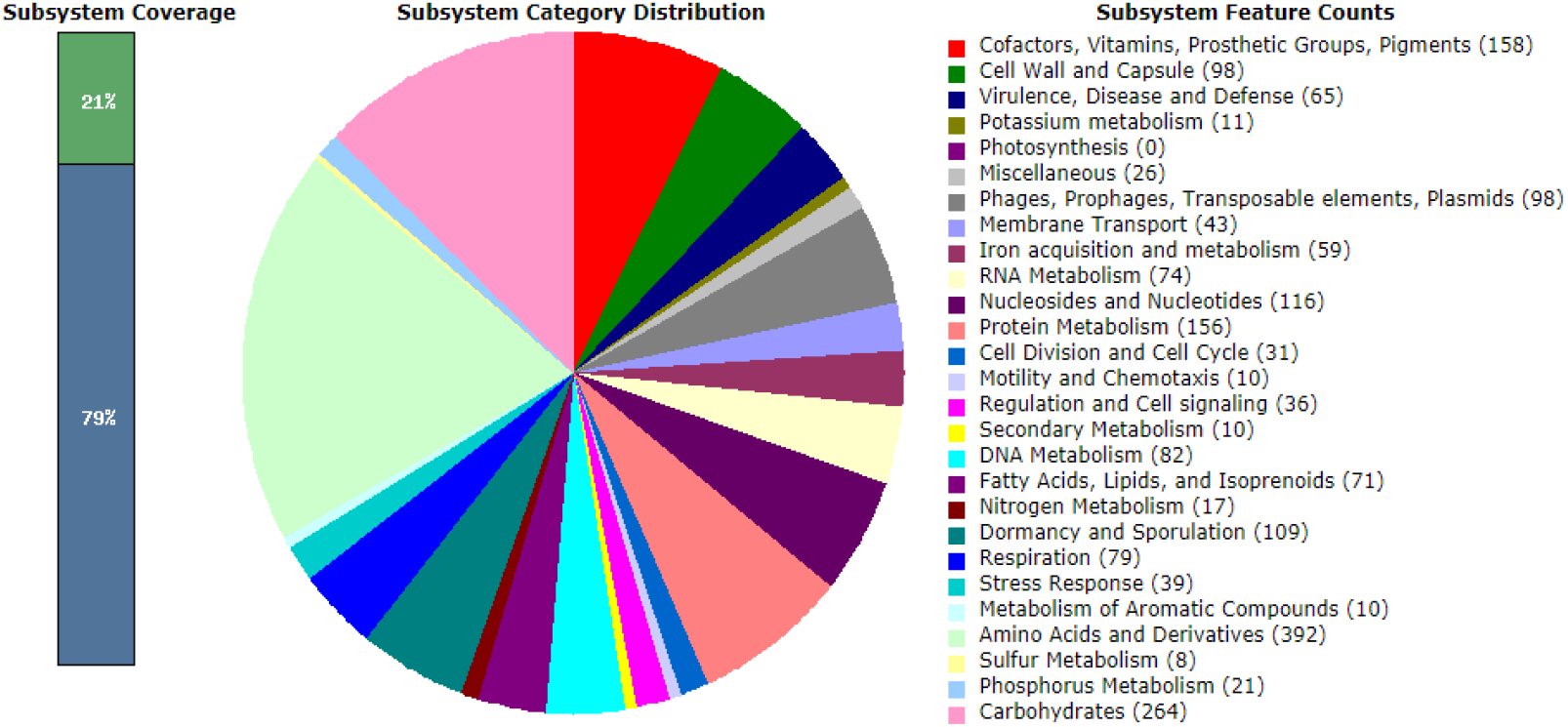
Functional annotation of the *Bt* SY49.1 strain genome sequence

**Figure 4.**
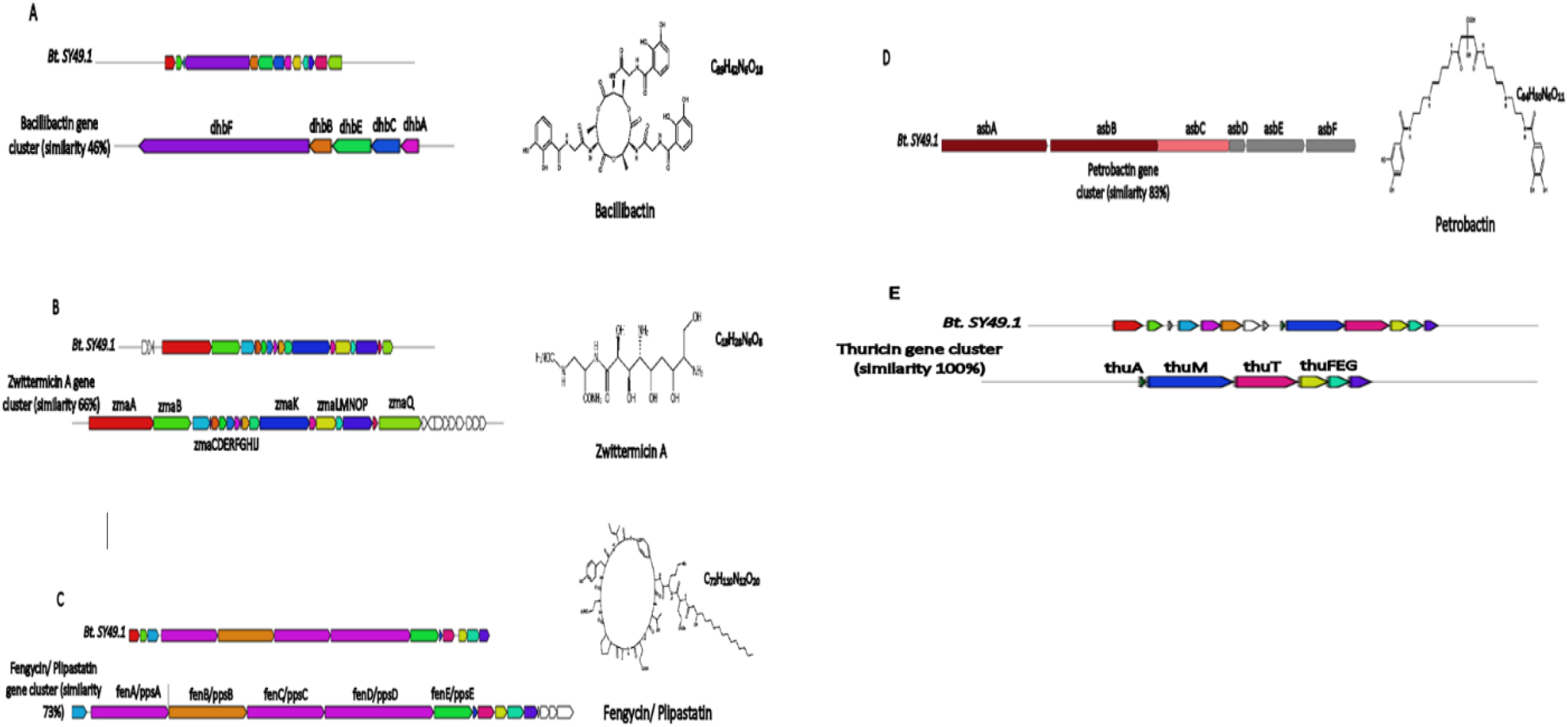
Secondary metabolite biosynthetic gene cluster organization in *Bt* SY49.1 genomes as predicted by the antiSMASH 4.0 server.

**Figure 5.**
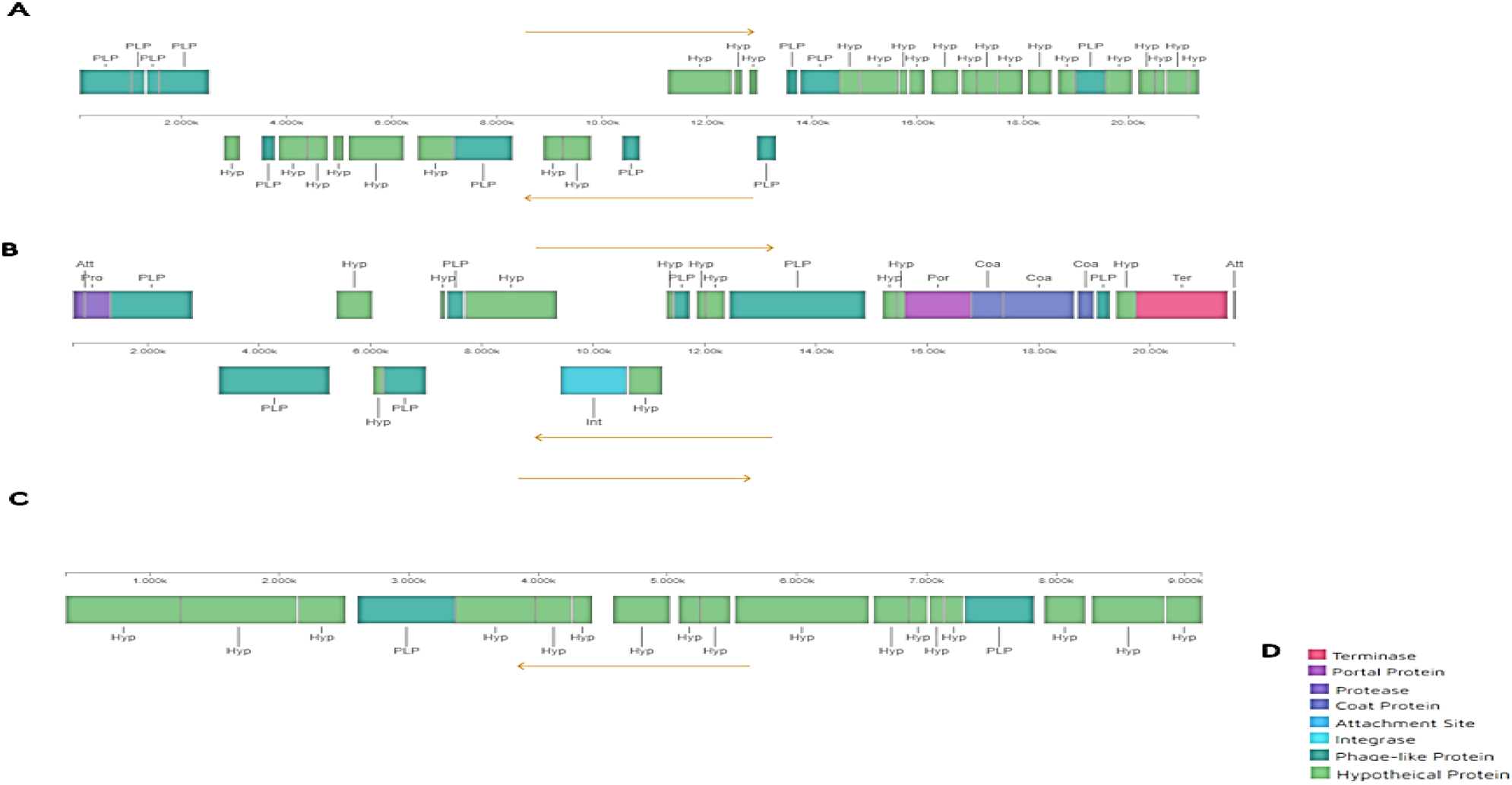
Prophage regions predicted in *Bt* SY49.1 genomes. 4A. Shows intact prophage region (score >90). 4B and 4C. show questionable prophage regions (score 70-90). 4D. shows the color legend.

### 3.3 Result of evolutionary and comparative genomic analyses

The phylogenetic tree of complete genome sequence of SY49.1 using PGFams and maximum likelihood tree of thirteen strains of *Bacillus* supports the placement of strain *Bt* SY49.1 within the *B. thuringiensis* group, see Figure 6. In addition, the genome of *Bt* SY49.1 is highly similar to *B. thuringiensis* serovar kurstaki strain HD 1 (CP010005) based on digital DNA: DNA Hybridization (DDH) estimate (GLM-based): 99.60%.

**Figure 6.**
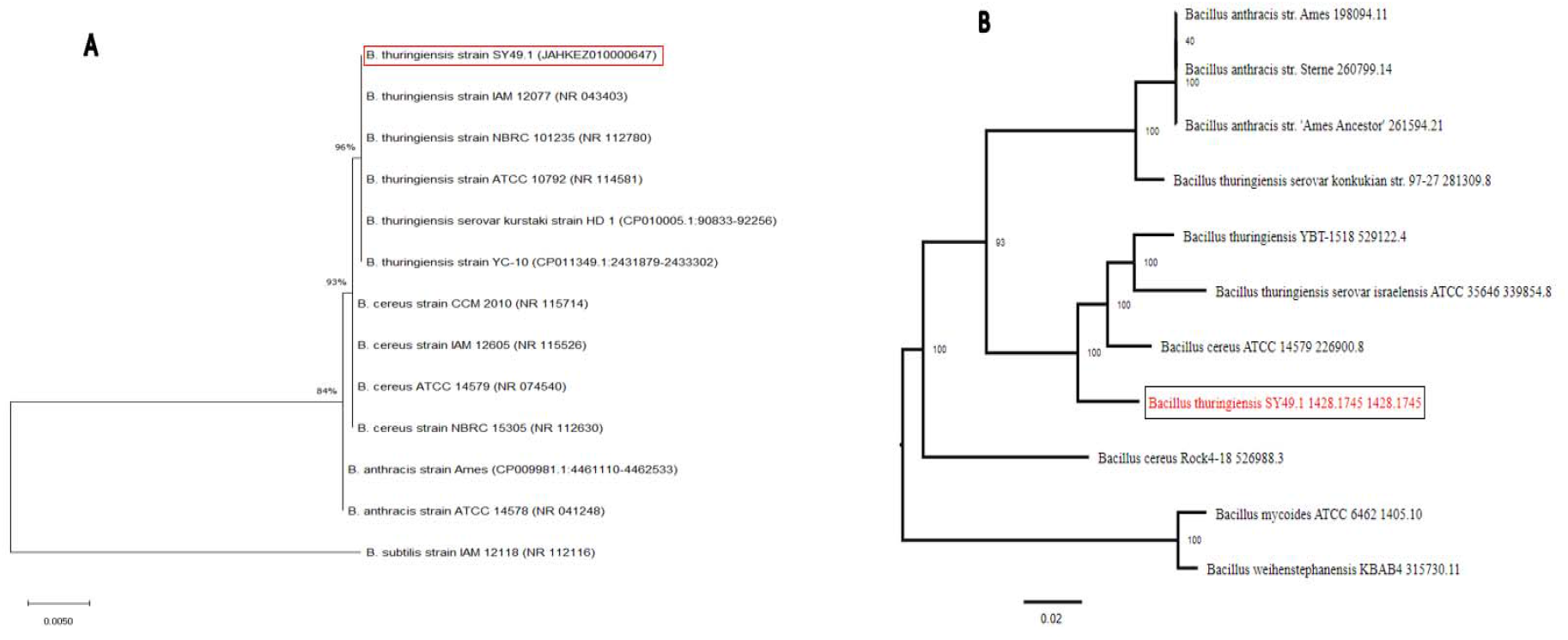
Molecular phylogenetic analysis of *Bt* SY49.1. The percentage of replicate trees (1000 replicates) is shown next to the branches. 5A. The maximum likelihood phylogenetic tree of *Bt* SY49.1 using *16S rRNA* sequences. The evolutionary distance was computed using the (K2+G) model. 5B. The phylogenetic tree of *Bt* SY49.1 complete genome using PATRIC global protein families (PGFams). The evolutionary analyses were conducted for *16S rRNA* and PGFams using MEGA7 and PATRIC software, respectively.

The Comparison of *Bt* SY49.1 genomes with the six closely related strains of *B. thuringiensis, B. cereus*, and *B. anthracis*, revealed strain-specific genes which encode hypothetical proteins, phage-like proteins, prophages, mobile genetic elements, and transposases. These findings were cross-validated and visualized in IslandViewer 4 server which showed these genes grouped into genomic islands, see Figure 7 and supplementary file 1: Table S1.

**Figure 7.**
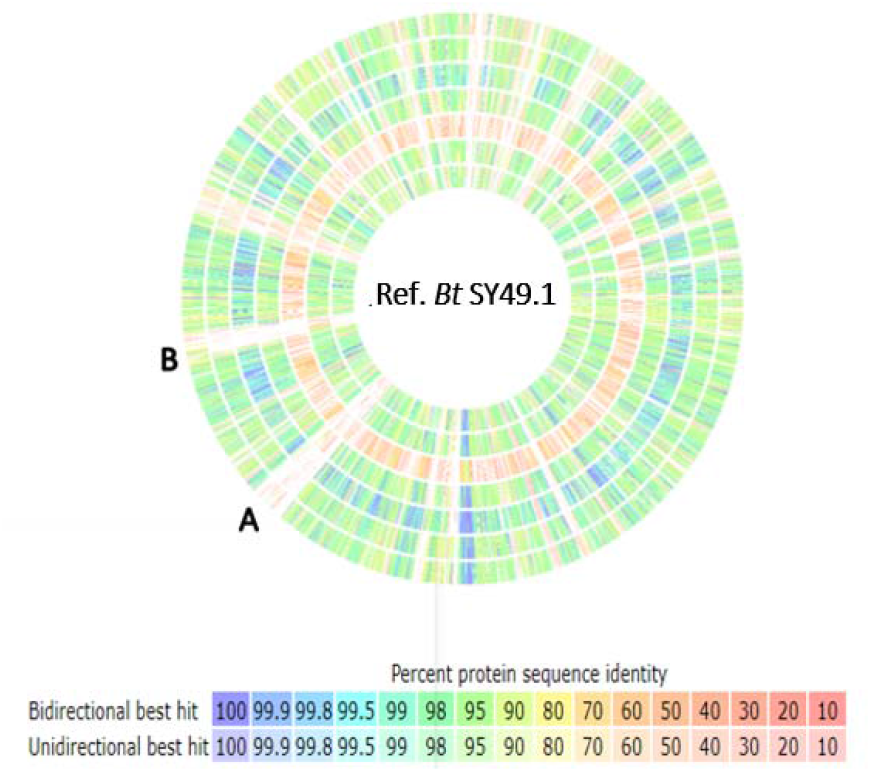
Genomic comparison of *Bt* SY49.1 to other *Bacillus* strains genomes. Each circle represents a pair-wise BLAST comparison between the open reading frames in query genome against those in *Bt* SY49.1 (Ref. = reference), with the percentage of similarity represented with different colors shown in the legend. Regions marked in the genomic map correspond to gene numbers presented in Additional File 1: Table S1, (A= 3993-4126) and (B= 4654-4792). Query genomes used in this analysis (outer ring to inner ring): *B. anthracis* str. Ames Ancestor (NC_007530), *B. cereus* ATCC 10987 (NC_003909), *B. cereus* ATCC 14579 (NC_004722), *B. cereus E33L* (NC_006274), *B. subtilis* subsp. *subtilis str. 168* (NC_000964), *B. thuringiensis* str. 97-27 (NC_005957), *B. thuringiensis* str. Al-Hakam (NC_008600).

Regarding variants calling, alignment of the *Bt* SY49.1 scaffolds to the reference sequence (NC_017208) revealed 122,508 SNPs and 160 InDels (6,206 bp), which accounts for 1.94% and 2.58%, respectively. Mutation types caused by InDels are presented in Table 1, and InDel statistics of their different size are shown in Figure 8. Among the 122,508 SNPs, 104,06 (84.94%) were in CDS region, while 18, 45 (15.06%) were identified in intergenic region. The distribution of SNPs in the CDS region is illustrated in Table 1.

**Table 1.** Mutation types cause by InDels

| Type | Frame shifted | Damaged start codon | Damaged stop codon | Premature stop | Other changes | CDS with InDel | Total CDS |
| --- | --- | --- | --- | --- | --- | --- | --- |
| No. (#) | 48 | 5 | 26 | 37 | 44 | 160 | 6,206 |
| Rate (%) | 0.773 | 0.081 | 0.419 | 0.596 | 0.709 | 2.578 | 100 |

Frame shifted, InDels induce the open reading frameshift mutation; Damaged Start Codon, the start codon is damaged by InDels; Damaged Stop Codon, the stop codon is damaged by InDels; Premature Stop, frameshifted cause a premature stop codon to occur, shortening the actual length by InDels; Other Change, there is no significant influence for the open reading frame with InDels.

**Table 2.**
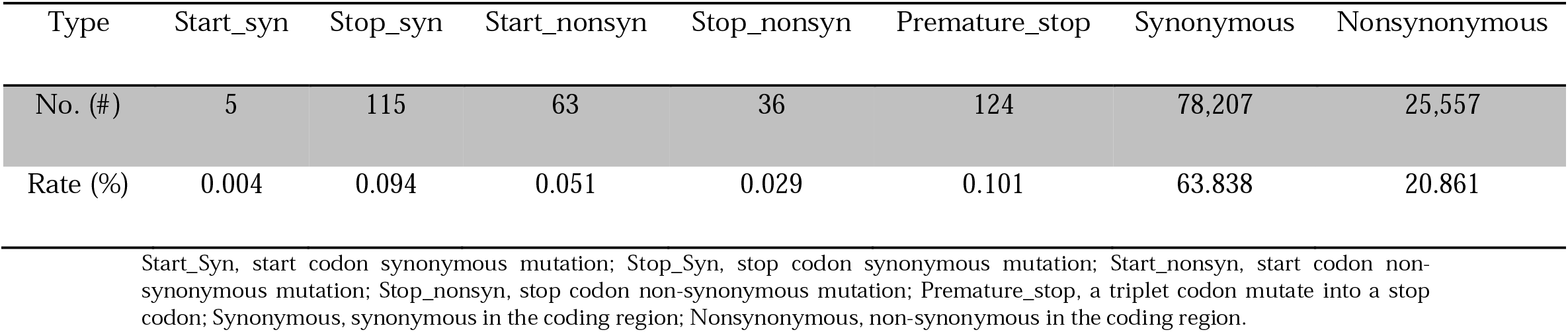
Distribution of SNPs in CDS region

**Figure 8.**
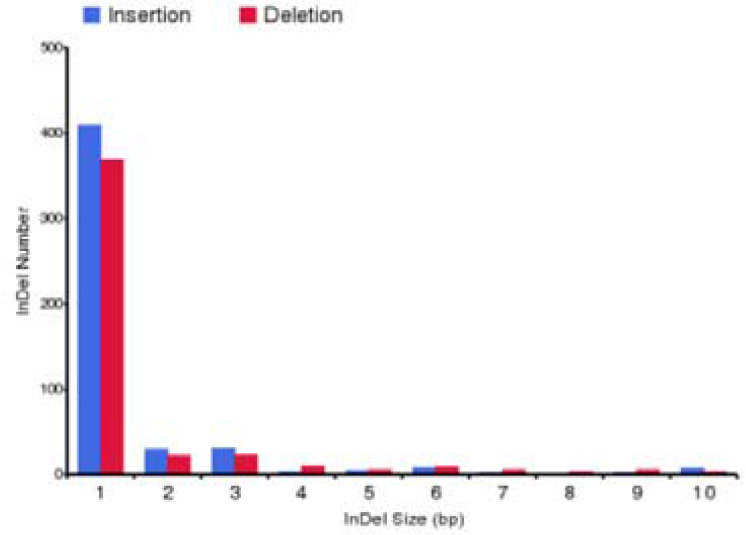
InDel statistics of different size

## 4. Discussion

In this study, we provided a standard genome draft of strain *Bt* SY49.1 which may contribute to the evolution and comparative genomics studies among the *B. cereus sensu lato* group. The draft genome of the *Bt* SY49.1 strain is 6.32 Mbp with 34.68% GC content, similar to the genomes of other *B. thuringiensis* strains (15)(16). In addition, the comparative analysis of strain *Bt* SY49.1 with closely related *Bacillus* genomes, the phylogenetic tree of the 16S rRNA sequences, and detection of gene inventories for the insecticidal Cry proteins biosynthesis placed *Bt* SY49.1 under *B. thuringiensis*. However, the *Bt* SY49.1 strain was selected for complete genome sequencing, annotation, and analysis as it was determined to be one of the most promising strains for the discovery of compounds with strong antibacterial, antifungal, and iron acquisition abilities (Figure 4).

Natural compounds produced by microorganisms form the basis of many drugs such as antibiotics, antivirals, and others (49). However, the rapid emergence of drug-resistant to recent antimicrobials and quick evolution through genetic alteration are great threats to control microbial infection (50). Therefore, research strategies like applying naturally occurring antimicrobial peptides (AMPs), and structural modification of antimicrobials drugs are extensively studied (51, 52). In this study, the *Bt* SY49.1 genome was found to carry gene clusters including lantibiotics, antimicrobial peptides, and siderophores. The antifungal compounds (fengycin and zwittermycin A) were found in the *Bt* SY49.1 genome. Fengycin, like most natural AMPs, acts by increasing the plasma membrane permeability of the target cell and exhibits strong fungi toxic activity (53, 54). Also, the aminopolyol antifungal compound (zwittermycin A) was previously shown to suppress oomycete diseases in plants (55, 56). These findings indicate that the *Bt* SY49.1 has antifungal activity besides chitinase, as indicated previously (24). Therefore, further experimental studies are also required to investigate the effectiveness of these compounds (fengycin and zwittermycin A) in different fungal diseases of plants. In addition, the *Bt* SY49.1 genome carries gene clusters of siderophores, like petrobactin and bacillibactin, which suggests its iron acquisition abilities. However, these gene clusters are not exclusive to *B. thuringiensis* genomes but are also detected in the genomes of other members of the *Bacillus cereus* group and other *Bacillus* species (57, 58).

Lantibiotics are gene-encoded peptide antibiotics that imply great potential to be attractive options for medical, agricultural, and industrial applications. Because of their special mode of action and they don’t raise substantial resistance. Thus, the discovery or detection of lantibiotics is urgent and appealing (59). However, the *Bt* SY49.1 genome was found to carry gene clusters with complete homology to the biosynthetic gene cluster of the thuricin (class II lantibiotic), see Figure 4E. The antimicrobial assay of thuricin exhibited its activity against *Micrococcus flavus*, and the conserved disulfide bridge was significant to the function of this bovicin HJ50-like lantibiotic (60). Interestingly, the antiSMASH 4.0 server predicted a number of novel secondary metabolites mainly belonging to the non-ribosomal peptide synthetase (NRPS) cluster and unspecified ribosomally synthesized and post-translationally modified peptide product (RiPP) cluster. Therefore, future investigations on *Bt* SY49.1 genome-encoded bioactive metabolites may be pursued.

## Conclusion

In this study, we report a 6,32 Mbp draft genome with 34.68% GC content of strain *Bt* SY49.1 isolated from the soil sample in Adana, Turkey. The assembled genome contains 6,562 protein-encoding genes, of which the most abundant are genes that are associated with amino acids and derivatives metabolism 392 (18.8%), followed by carbohydrates metabolisms 264(12.67%), and cofactors, pigment, and prosthetic groups 158(7.59%). The antibacterial, antifungal, and insecticide properties of this strain could be inferred, in part, with several gene inventories encoded in the draft genome. This indicated that strain *Bt* SY49.1 could have several potential utilities as a source of antibiotics compounds, and a fungal phytopathogen and insect biocontrol agent. We expect that the draft genome of the *Bt* SY49.1 strain may provide a model for proper understanding and studying of antimicrobial compound mining, genetic diversity among the *Bacillus cereus* group, and pathogenicity against insect pests and plant diseases.

## Data availability

The *Bt* SY49.1 whole-genome sequencing project has been deposited at NCBI GenBank under the accession number NZ_JAHKEZ000000000, BioProject accession number PRJNA734785, and BioSample accession number SAMN19533886.

## Acknowledgments

This work was supported by Erciyes University Scientific Project Unit under the codes of ÖNAP-3638 and FBD-08-540. We also thank to GenKök research center for their collaboration.

